# In-depth evaluation of root infection systems with the vascular fungus *Verticillium longisporum* as soil-borne model pathogen

**DOI:** 10.1101/2020.12.28.424556

**Authors:** Christian Fröschel

## Abstract

**PREMISE:** While leaves are far more accessible for analysing plant defences, roots are hidden in the soil leading to difficulties in studying soil-borne interactions. Literature describes inoculation strategies to infect model plants with model root pathogens, but it remains demanding to obtain a methodological overview. To address this challenge, this study uses the model root pathogen *Verticillium longisporum* on *Arabidopsis thaliana* and provides recommendations based on evident examples for the selection and management of suitable infection systems to investigate root-microbe interactions.

**METHODS AND RESULTS:** A novel root infection system is introduced, while two existing ones are precisely described and optimized. Advantages and disadvantages of each are assessed, step-by-step protocols are presented and accompanied by pathogenicity tests, transcriptional analyses of indole-glucosinolate markers and independent confirmations using reporter constructs. The results validate the importance of indole-glucosinolates as secondary metabolites limiting *V. longisporum* propagation in hosts.

**DISCUSSION:** We provide detailed guidelines for studying host responses and defence strategies against *V. longisporum*. Furthermore, other soil-borne microorganisms or other model plants, such as economically important oilseed rape, can be used in the infection systems described. Hence, these proven manuals help to find a root infection system for your specific research questions to decipher root-microbe interactions.

## INTRODUCTION

Soil-borne microorganisms affect plant growth and can cause disease. Within the fungal genus *Verticillium (Ascomycota*) some members are well-known plant pathogens that cause so-called vascular wilt diseases, leading to an estimated annual loss of €3 billion worldwide (Depotter *et al*., 2016). One important agent is *Verticillium longisporum*, which is highly adapted to brassicaceous hosts, such as oilseed rape (*Brassica napus*) and cauliflower (Depotter *et al*., 2016). Nowadays, oilseed rape is one of the top three sources of vegetable oil in the world and of economic importance by providing human food, animal feed and biofuels (Bancroft *et al*., 2011, Carré *et al*., 2014, USDA, 2016). Depending on environmental conditions, yield losses in oilseed rape cultures caused by *V. longisporum* might locally range between 10 % and 50 % (Daebeler *et al*., 1988; Dunker *et al*., 2008, Depotter *et al*., 2016). The infection process initiates in the soil from resting forms, which germinate in response to plant exudates (Mol and Scholte, 1995). The fungal hyphae traverse the outer root cell-layers by intra- and intercellular growth and penetrate the vasculature in the inner central cylinder (Eynck *et al*., 2007, Depotter *et al*., 2016). Most of its life cycle, the fungus exists in the vascular tissue. In there, the xylem sap is nutrient-poor and delivers plant defence compounds, so that *Verticillium* spp. have to adapt to this unique environment by a fine-tuned secretion of colonization-related proteins enabling the pathogen to react to changes in nutrient supply or host defences (Floerl *et al*., 2008; Singh *et al*., 2010; Leonhard *et al*., 2020). This adaption allows the fungus to grow. Once in the vasculature, conidiospores are released and disseminated acropetally with the water stream, leading to a systemic spread in the host and colonization of the foliage (Depotter *et al*., 2016; Iven *et al*., 2012). By then, the fungus affects plant growth (Eynck *et al*., 2007; Floerl *et al*., 2008; Fröschel *et al*., 2019), triggers premature senescence (Reusche *et al*., 2013) and since the infection progress impairs plant’s water transport, typical symptoms like wilting, stunting and yellow leaves occur (Zhou *et al*., 2006). Thus, crop yield and quality of infected plants is strongly reduced. As up to date no efficient fungicides are available (Depotter *et al*., 2016), further research is required to define genetic determinants of resistance to develop novel strategies for disease control in agriculture.

In response to pathogens, plants evolved inducible defence mechanisms. As one efficient strategy, plants produce secondary metabolites to defend themselves against invading microorganisms (e.g. Ahuja *et al*., 2012). The so-called tryptophan-derived secondary metabolites arise from the amino acid tryptophan. Among these, sulfur-containing camalexin and indole-glucosinolates (IG) are described in the model plant *Arabidopsis* to combat various plant pathogens (e.g. Clay *et al*., 2009; Bednarek *et al*., 2009). Modifications in the biosynthetic pathway of these compounds, such as hydroxylation and subsequent methylation, require several CYTOCHROME-P450 enzymes (CYPs) (reviewed in Halkier and Gershenzon, 2006). The key enzymes in the beginning of the biosynthesis are CYP79B2 and CYP79B3. The double mutant *cyp79b2/b3* shows increased susceptibility to a variety of microorganisms, such as *Phytophthora brassicae* or *V. longisporum* (Schlaeppi *et al*., 2010; Iven *et al*., 2012), attesting the importance of these secondary metabolites for defence. In *V. longisporum* infected *Arabidopsis* roots, gene expression of the IG pathway is heavily induced (Iven *et al*., 2009, Fröschel *et al*., 2019). A prominent example is the strong induction of *CYP81F2*, which encodes an important enzyme in the terminal part of IG biosynthesis. Recently, some transcription factors from the ETHYLENE-RESPONSE-FACTOR (ERF) family were shown to be activators of genes necessary for IG production (Xu *et al*., 2016). Overexpression of several closely related ERFs enhances resistance against *V. longisporum*, such as ERF105, which directly binds to the *CYP81F2* promoter (Fröschel *et al*., 2019).

To investigate interactions of roots with soil-borne microorganisms, several studies combine *Arabidopsis thaliana* and *V. longisporum* (Depotter *et al*., 2016). Central reasons for using these models are easy-to-handle susceptibility assays and the well-established genetic resources to study expressional alterations. Through comparative analysis of the molecular events explained above, we optimized two existing infection systems and developed one new *in vitro* infection system. We highlight advantages and disadvantages of each infection system helping to find an appropriate application for specific research questions in general root-microbe interactions.

## METHODS

### Material

The virulent *Verticillium longisporum* (*Vl*) strain *Vl*43 was originally isolated from *Brassica napus* in northern Germany (Zeise and von Tiedemann, 2002a, b). As this strain was frequently used in publications, it was also taken here. *Vl*43 possess cefotaxim resistance. *Vl*-sGFP (i.e. *Vl*43 constitutively expressing enhanced Green Fluorescent Protein; Eynck *et al*., 2007) was used as fluorescent fungal reporter line. *Arabidopsis thaliana* wild type (WT) and all transgenic lines had Col-0 background: *cyp79b2/b3* (Zhao *et al*., 2002), *ERF105* overexpressing line (*ERF105 OE*) (Fröschel *et al*., 2019), *Promoter_CYP79B2_:LUC* (Iven *et al*., 2012). WT oilseed rape (*Brassica napus* var. *napus*) cultivar “Miniraps” (rapid-cycling rape genome ACaacc; Floerl *et al*., 2008; Williams and Hill 1986).

### Surface sterilization of seeds

Both seeds from *A. thaliana* and *B. napus* were surface sterilized by incubation with chlorine gas for 3 h prior usage in the infection systems. Chlorine gas was generated in a desiccator containing the seeds through the reaction of 100 ml of 12% sodium hypochlorite (NaClO in H_2_O) and 6 ml of 33 % hydrochloric acid (HCl).

### Cultivation of V. longisporum and inoculum preparation

Note, all equipment and medium had to be sterile. Additionally, all steps were carried out in a laminar flow hood to keep the inoculum clean. To obtain mitotically derived (asexual) conidiospores, mycelia had to be cultivated in liquid Potato Dextrose Broth medium (PDB; Sigma-Aldrich) supplemented with 500 mg/l cefotaxim (Duchefa). A 500 ml chicanery flask was filled with 150 ml PDB medium and inoculated with *Vl*43 spores from a glycerol-stock storage. 7-10 days incubation in darkness at room temperature (RT) under continuous, horizontal shaking (rotary shaker, 60 rpm) resulted in small, spherical, white mycelia (Appendix S1A, B). The PDB medium was carefully removed, while the mycelia remained in the flask. To induce sporulation, 100 ml of Czapek Dextrose Broth medium (CDB, Duchefa) supplemented with 500 mg/l cefotaxim was added to the mycelia. Another 4-5 days incubation in darkness at RT under continuous, horizontal shaking (rotary shaker, 60 rpm) resulted in a yellow-greyish supernatant (Appendix S1C) containing the conidiospores. To separate the spores from the mycelia, a portion of the supernatant was filtered through filter paper (pleated cellulose filter, particle retention level 8-12 μm) into a sterile 50 ml collection tube (Appendix 1D, E). By using a Thoma cell counting chamber, the spore-concentration was determined and diluted with sterilized ¼ MS (Murashige & Skoog medium including vitamins, Duchefa) in Milli-Q water until the final spore-concentration was achieved (details see below). Freshly harvested conidiospores were always used as inoculum ^**1)**^. For long term storage, isolates were frozen as high concentrated spore solutions (approx. 1×10^8^ spores/ml) in 25 % glycerol at − 80 °C. 100 μl of those glycerol stocks were used to inoculate the PDB medium in the beginning of this section.

### Sterile in vitro infection system in petri dishes

The petri dish cultivation system based on a medium consisting of 1.5 g/l MS (Murashige & Skoog medium including vitamins, Duchefa) and 8 g/l Gelrite Agar (Carl Roth GmbH) in Milli-Q water ^**2)**^. After autoclaving (121 °C), the medium was poured into petri dishes (92 x 16 mm, Sarstedt AG & Co. KG). All steps were carried out in a laminar flow hood with sterile equipment. The upper third as well as an infection channel were cut and removed with a scalpel from the solidified medium ^**3)**^ as illustrated in Appendix S1F. Per plate, 50-100 surface sterilized *Arabidopsis* seeds were distributed with a pipette tip on the cut surface directly at the angle to the petri dish wall (allowed roots to grow between medium and petri dish wall, which facilitated inoculation and harvesting compared to a root growth completely in medium). The petri dishes were sealed with Leukopor^®^ (BSN medical GmbH) allowing gas exchange. After stratification for 2 d in darkness at 4 °C, the plates were set up vertically and plants were grown at 22 °C ± 1 °C under long day conditions (16 h light / 8 h darkness) in a poly klima^®^ growth chamber (poly klima GmbH). As soon as the roots reached the channel (9-11 d old seedlings), 500 μl of freshly harvested conidiospores (4×10^5^ spores/ml) were added to the infection channel (Appendix S1G). To prepare control plates, 500 μl mock solution without spores was used. After adding the solutions, the plates were incubated horizontally in the hood for at least 40 min before they were sealed with Leukopor^®^. The root parts were covered with black paper-boxes and plates were placed vertically again (Appendix S1H). To control success of infection, it was demonstrated by microscopy that infected roots differed from mock treated ones (Appendix S1I). At the time points indicated, leaves were cut from the roots and harvested separately. By taking out the agar strips from the petri dishes, roots became easily accessible and were carefully pulled out of the agar using forceps. All material was flash-frozen in liquid nitrogen and used either for analyses of transcriptional changes or determination of fungal DNA. Each biological replicate consisted of pooled leaves or roots from two plates.

### Soil-based infection system in pots

This system was kept semi-sterile. A 3:1 (v/v) soil:sand mixture (bird sand from Pet Bistro, Müller Holding Ltd. & Co. KG) was steamed ^**4)**^ and filled in pots (∅ 7 cm). The pots were transferred to trays and water was added into the trays about 1/3 the height of a pot. 3 h later the soil was thoroughly soaked with water. In addition, the soil surface was water-sprayed with a spray bottle, to ensure wet soil conditions. 3-4 seeds were sawn per pot and stratified for 3 d in darkness at 4 °C to synchronize germination. Thereafter, plants were cultivated for 21 days under long-day conditions (16 h light / 8 h darkness; 22°C; 60 % humidity) and regular watering ^**5)**^. To reduce individual variations, an excess of plants was pre-cultivated and just plants of similar size were carefully selected for the actual experiments. Appendix S2A and B illustrate the procedure, called “root dip inoculation” (Iven *et al*., 2012). 21 d old *Arabidopsis* plants were prepared: After pulling the soil out of the pots, roots were carefully excavated, gently washed in a water container ^**6)**^ and dipped for 60 min in a petri dish holding 35 ml of spore-solution (2 x 10^6^ spores/ml) or a mock solution without spores. Following the 60 min incubation, plants were inserted ^**7)**^ into single pots containing moist, steam-sterilized soil ^**4)**^ without sand and cultivated under long-day conditions (16 h light / 8 h darkness; 22°C; 60 % humidity) with regular watering ^**5)**^. Solely the rosettes were harvested by cutting them at the root crown at the time points indicated. To analyse fungal DNA, rosettes of 5 plants were pooled per replicate and at least 3 biological replicates were analysed per line. All material was flash-frozen in liquid nitrogen.

### Sterile in vitro infection system in plastic cups

A novel sterile *in vitro* system in plastic cups was established. All steps were carried out in a laminar flow hood with sterile equipment. All plastic ware was sterilized in a 70 % ethanol bath for at least 15 min. Appendix S3 illustrates the principle of this infection system. In brief, a MS (Murashige & Skoog medium including vitamins, Duchefa) based Gelrite Agar (Carl Roth GmbH) medium ^**2)**^ (4.4 g/l MS, 0.2 g/l MgSO_4_, 1 g/l KNO_3_, 0.5 g/l MES [2-(*N*-morpholino)ethanesulfonic acid], 6.0 g/l Gelrite in Milli-Q water, adjust pH 5.7 with KOH) was autoclaved (121 °C) and poured into transparent 500 ml plastic cups (108 x 82 x 99 mm, salad boxx^®^ Feinkostbecher) ^**8) 9)**^. A separating plastic layer was transferred to the medium before it solidified. It contained holes for placing surface sterilized seeds onto the medium. This allowed the seeds an access to the medium and later it prevented the leaves from touching the fungus containing medium. Therefore, all fungal DNA detectable in leaves derived solely from fungal spread within the plant. As soon as the medium solidified, seeds were transferred to the prefabricated holes ^**10)**^ and an approx. 1.5 cm deep infection channel was cut through a centre hole, which allowed to add fungal spores later. The cups were covered with a second, inverted plastic cup and sealed with Leukopor^®^ (BSN medical GmbH) ^**11)**^. To synchronize germination, seeds were stratified for 3 d in darkness at 4 °C. Seedlings were grown under 12 h light / 12 h darkness conditions in percival chambers (CLF Plant Climatics GmbH) at 22 °C and 60 % humidity. In case of *Arabidopsis*, 21 d old seedlings were inoculated by adding 1 ml conidiospore solution (4×10^5^ spores/ml) into the infection channel cut in the beginning. For control samples, 1 ml mock solution without spores was added. The whole rosettes were harvested at time points indicated and prepared to analyse the amount of fungal DNA or expressional changes. Roots were carefully pulled out of the medium, squeezed and dabbed with a paper towel to remove agar residues. Each replicate consisted of rosettes / roots pooled from 5 plants and at least 3-4 biological replicates were analysed per line or time-point. All material was flash-frozen in liquid nitrogen. *B. napus* was similarly treated like *Arabidopsis*, but as this species grows faster, 5-7 d old seedlings were inoculated. At the time points indicated, all leaves present on each *B. napus* seedling were harvested and leaves from 5 plants were pooled for each replicate. All *B. napus* material was flash-frozen in liquid nitrogen to determine fungal DNA.

### Quantification of V. longisporum DNA via quantitative PCR (qPCR)

The amount of *V. longisporum* DNA was quantified in leaves to estimate fungal propagation *in planta*. After grinding the leaf material in liquid nitrogen, 100 mg powder was transferred to a 2 ml reaction tube and used to extract total DNA (CTAB [cetyltrimethylammonium bromide] method for DNA extraction modified from Clarke, 2009: 400 μl CTAB extraction buffer was added to each sample, followed by vortex and a 15 min incubation at 65 °C. 400 μl chloroform:isoamyl alcohol (24:1, v/v) was added, followed by vortex and centrifugation at 13,000 rpm for 5 min at RT. 280 μl of the aqueous layer was transferred to a clean tube and 280 μl isopropanol was added. It was mixed by inversions and incubated for 3 min at RT. It was centrifuged at 13,000 rpm for 10 min at RT. The supernatant was discarded and the pellet was washed with 300 μl cold 70 % ethanol. A brief centrifugation secured the pellet so that the supernatant could be carefully removed with pipettes. The pellet was dried and the DNA was dissolved in water. The concentration was adjusted to approx. 50 ng/μl). qPCR was performed with 100 ng total DNA and BIOTAQ™ DNA polymerase (Bioline GmbH) in a CFX96™ Real-Time PCR Detection System (Bio-Rad Laboratories, Inc.). Amplification products were visualized by SYBR^®^ Green (Lonza Group AG). Following conditions were generally applied: 10 min at 95°C, 40 cycles of: 20 s at 95°C, 20 s at 56°C and 20 s at 72°C, followed by a quality check program determining the melting curve to detect nonspecific amplification. Fungal DNA was determined by *Verticillium* specific primers OLG70 and OLG71 (Eynck *et al*., 2007) and evaluated relative to an *ACTIN8 (ACT8*) amplicon derived from *A. thaliana* or *B. napus* genomic DNA (Fröschel *et al*., 2019). Quality control was performed showing specificity of the primers depending on DNA type (Appendix S4). Relative amount of fungal DNA was calculated as x-fold over WT or mock by 2^−ΔΔCT^ method (Livak and Schmittgen, 2001). One data point was calculated from 3-5 biological replicates (*n*). Primer sequences are given in Appendix S5. Determining fungal DNA from root material is difficult, as it cannot be discriminated between mycelia that have grown outside or inside the roots, which would be a requirement to analyse host susceptibility.

### Detection of transcriptional responses via qRT-PCR

Each RNA sample was isolated from roots or leaves following the TRI Reagent procedure (Chomczynski and Mackey 1995). 30 min of DNase I treatment (Thermo Fisher Scientific, Inc.) was performed to eliminate DNA contaminations in RNA extracts. cDNA was synthesized from 1 μg of total RNA using oligo(dT) primers, random nonamer primers and the reverse transcriptase RevertAid H Minus (Thermo Fisher Scientific, Inc.) according to the manufacturer’s manual. 2 μl of 1:10 diluted cDNA and BIOTAQ™ DNA polymerase (Bioline GmbH) were used in a CFX96™ Real-Time PCR Detection System (Bio-Rad Laboratories, Inc.). Amplification products were visualized by SYBR^®^ Green (Lonza Group AG). The following conditions were generally applied: 10 min at 95°C, 40 cycles of 20 s at 95°C, 20 s at 56°C and 20 s at 72°C, followed by a default dissociation stage program to detect nonspecific amplification. Primers are given in Appendix S5. Values were calculated from at least 3 biological replicates as fold induction values using 2^−ΔΔCT^ method (Livak and Schmittgen 2001). *UBQ5* served as reference gene for normalization.

### Analysis of green leaf area and fresh weight

After removing the stems, photographs of the plants were taken with a digital camera always at the same distance from above. Projected green leaf area was assessed by using *BlattFlaeche^®^* (Datinf GmbH, Tübingen, Germany) software as described in Iven *et al*., 2012 and Appendix S6A. The diameter of the culture pots was used as an internal standard for normalizing the leaf area. Biomass (fresh weight) of *Arabidopsis* rosettes was determined after removing roots and stems by weighing whole rosettes. Relative fresh weight was determined by normalizing fresh weight of infected samples to that of the corresponding mock samples.

### Analysis of fungal colonization in stems

21-30 dpi (days post inoculation), 1–1.5 cm of primary inflorescence stem segments was cut at the base (Appendix S6B), surface sterilized (5 min, 70 % ethanol; 5 min 0.02 % sodium hydrogen chloride solution supplemented with 0.02 % Triton X-100) and thoroughly washed with sterile water (according to Fröschel *et al*., 2019). These stem segments were placed on solidified Potato Dextrose Broth agar (20 g/l PDB (Sigma-Aldrich); 10 g/l phyto agar (Duchefa); Milli-Q water; autoclaved at 121 °C; supplemented after cooling to 60 °C with 500 mg/l cefotaxim (Duchefa)). After incubation in darkness for 3-5 d at RT, fungal mycelia became visible.

### Fluorescence confocal microscopy

*Arabidopsis* was inoculated in the petri dish system with *Vl*-sGFP conidiospores. Roots were transferred to microscope slides, water incubated and covered with coverslips. Examination was achieved using Leica TCS SP5 II confocal laser scanning microscope (Leica Camera AG). Samples were excited by Argon Ion Laser and the fluorescence signal for GFP was detected (excitation wavelength: 488 nm; detection window: 500-530 nm). Pictures were taken with a HC PL APO 20×0.70 IMM CORR CS water immersion objective. For propidium iodide (PI) staining, roots were incubated in darkness for 10 min in 15 μM PI (P4170, Sigma-Aldrich). Thereafter, roots were transferred to microscope slides, incubated in water and carefully covered with coverslips. PI fluorescence signal was detected (excitation 488 nm, detection window 610–670 nm) and merged with GFP channel.

### Luciferase assay

*Arabidopsis* reporter line *Promoter_CYP79B2_:LUC* was inoculated with *V. longisporum* in the system in petri dishes. Roots were sprayed 3 dpi with Luciferin solution (1 mM Luciferin, 0.02% Triton-X 100, demineralized H_2_O). Plates were incubated in a dark box of the luciferase imaging system C4742-98 coupled with cooled CCD camera (Hamamatsu Photonics K.K.). Black/white and luminescence pictures were taken. Intensities of luminescence were given in false colour (low intensity in blue; high intensity in red).

### Statistical analysis

Calculations and graphs were made with Microsoft^®^ Excel software. Statistical significance was assessed by using two-tailed Student’s *t*-test. Differences between two groups were considered significant with * *p ≤* 0.05, ** *p ≤* 0.01 or *** *p ≤* 0.001. Error bars indicate ± SD (standard deviation) or ± SEM (standard error of the mean) as indicated in the figures.

### Gene accessions

*ACTIN8* (At1g49240); *CYP79B2* (At4g39950); *CYP79B3* (At2g22330); *CYP81F2* (At5g57220); *ERF105* (At5g51190); *UBQ5* (At3g62250).

### Notes

1. Always use freshly harvested conidiospores to prepare solutions for experiments and not frozen storage stocks. Storage stocks lose germinability and viability, affecting reproducibility in independent experiments.
2. Avoid sugar in the agar medium as it will lead to excessive fungal growth after inoculation.
3. Keep the solidified petri dishes upside down in the fridge (4-10 °C) overnight before cutting. A chilled medium helps to prevent sliding after cutting. Avoid getting liquid or air under the agar medium while cutting, to circumvent sliding and to maintain the infection channel.
4. Do not autoclave at 121 °C as this will affect important organic compounds in the nutrition soil. Steaming at 80-100 °C for 20 min in an autoclave is fine to reduce microbial contaminations in the substrate to a very low rate.
5. Since *Arabidopsis* is sensitive to flooding stress, water only when there is no more standing water in the tray.
6. It is important that you bathe and wash just the roots in water, while keeping the rosettes out of the water between your fingers. If rosettes are too waterlogged, it can cause additional side effects from hypoxic stress.
7. To facilitate pants’ repotting, a hole should be made in the soil. After inserting the roots, make sure to refill the holes very softly with soil and avoid pressing. Otherwise, stress symptoms can result from repotting (e.g., blueish leaves).
8. As alternative, for example Phytatray™ II (Sigma-Aldrich) or Magenta™ vessel (Sigma-Aldrich) can be used. Although they are more expensive, they are delivered ready-to-use sterile.
9. Try to make 10-15 % more cup systems as you would need, because it happens occasionally that some seedlings do not grow properly.
10. Slightly scratch the solidified agar medium in the holes before placing the seeds on it. This interrupts the solidified skin at the surface, allows the seeds to soak with water and facilitates the radicle to penetrate the agar medium.
11. As it was obligatory to ensure a slight gas exchange with the environment, Leukopor^^®^^ was used.

## RESULTS

Plant responses towards the fungus were compared using three different infection systems. In all cases, *Arabidopsis* roots were inoculated with *V. longisporum* and expression of prominent marker genes (*CYP79B2, CYP79B3, CYP81F2* and *ERF105*) was analysed by comparing mock treated and infected plant samples using qRT-PCRs. Susceptibility of relevant transgenic lines was classified by determining the amount of *V. longisporum* DNA in leaves with qPCRs. In general, IG are important to limit *V. longisporum* spread (Fröschel *et al*., 2019). Therefore, a mutant and an overexpressor line of this pathway served to test reliability of the systems in detecting differences in infection progress.

### The petri dish system is best suited to study early expressional changes

First, the system was described to analyse root transcriptional responses against *V. longisporum* in Iven *et al*., 2012. Here, we optimized the protocol. Soon after inoculation, fungal mycelia became visible all around the root and in the medium (Fig. 1A). Fungal hyphae appeared all around the root tip and penetration of outer root tissue was observable using a GFP-tagged *V. longisporum* strain (*Vl*-sGFP) (Fig. 1B). Fungal DNA was detectable in leaves as early as 1 day post inoculation (dpi) at roots, which indicates a rapid systemic spread *in planta* in this infection system (time course in Fig. 1C). An almost exponential fungal propagation occurred until 3 dpi, followed by a flattening of the curve. In roots, induction of IG marker genes was very high with *CYP79B2, CYP79B3* and *CYP81F2* having their maximum value at 2 dpi and *ERF105* at 3 dpi (Fig. 1D). Beyond their induction maxima, expression decreased consecutively indicating an important role of IG especially in early infection. A reporter construct confirmed activation of the *CYP79B2* promoter in infected roots (Fig. 1E). Root and leave responses can easily be examined separately in this system. Induction of IG marker genes in leaves reached the maximum at 3 dpi or even later (Fig. 1F), thus time-delayed to root reactions. The knockout mutant *cyp79b2/b3* dramatically lacks IG (Zhao *et al*., 2002) and it was significantly more susceptible than the WT, whereas the *ERF105* overexpressor (OE) tended to be more tolerant in this infection system, although not significant (Fig. 1G). Overall, this system is well suited to examine early host responses both in roots and leaves by qRT-PCR and reporter gene constructs, but it has its limitations with categorizing susceptibility.

**Fig. 1:**
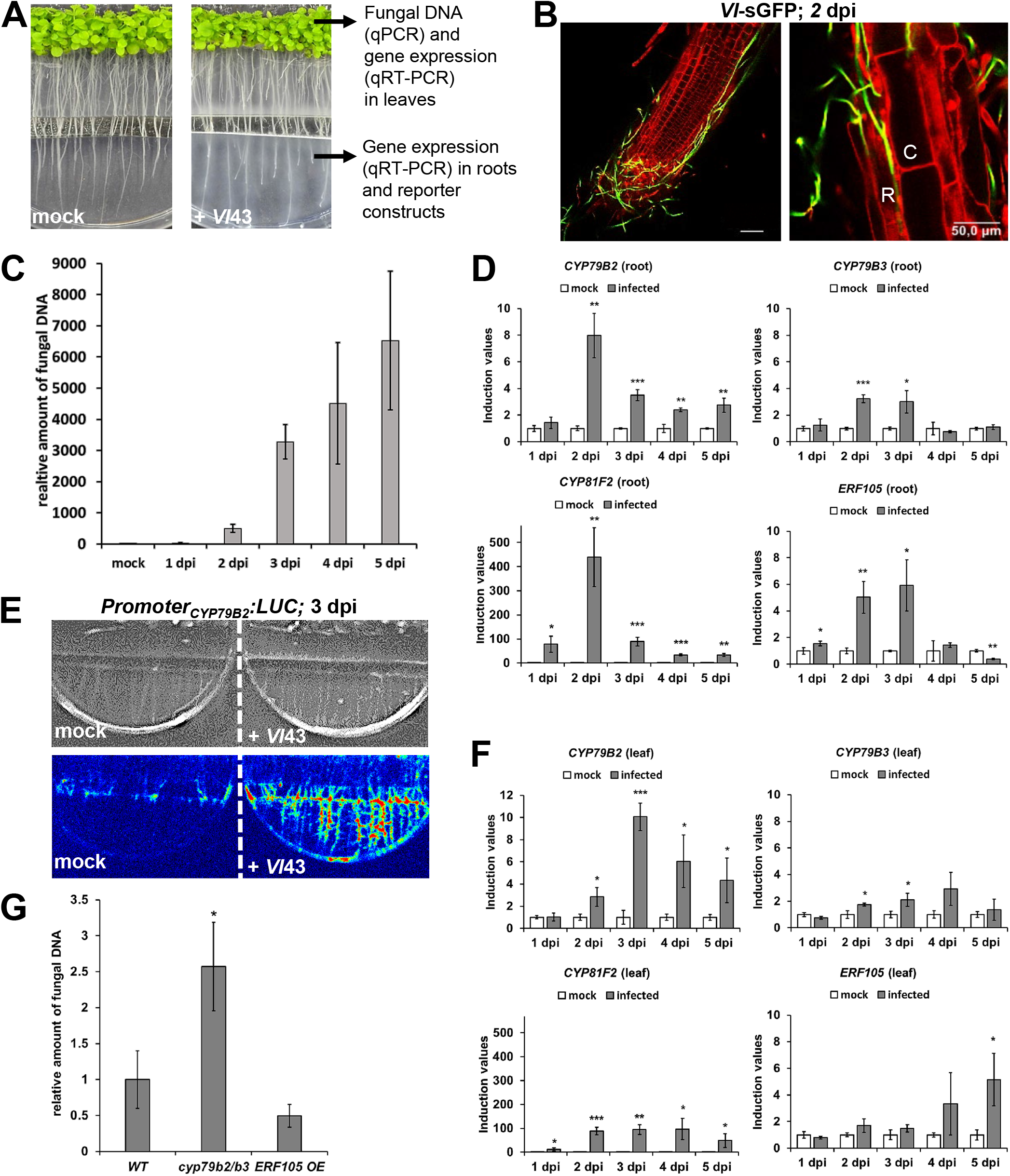
Infection system in petri dishes to study early *Arabidopsis-Verticillium* interactions. **A.** Visible growth of *V. longisporum* (*Vl*43) mycelium 3 days post inoculation (dpi). Representative pictures from inoculated (+ *Vl*43) and mock treated samples. Possibilities for downstream analyses are given. **B.** Visualization of fungal growth by using a GFP-tagged *V. longisporum* strain (green, 2 dpi). Counterstaining of root-cells with propidium iodide (red). A net of fungal hyphae around the root tip (left) and hyphal growth between cells of rhizodermis (R) and cortex (C) (right). Bars indicate 50 μm. **C.** Relative amount of *Vl*43 fungal DNA as quantified in WT leaves at the time points indicated. Values of infected samples are given relative to background noise in mock sample (set to 1) (*n* = 3 each, ± SD). **D.** Induction of marker genes in WT roots at the indicated time points. Values of *Vl*43 infected samples are given relative to mock samples (set to 1) (*n* = 3 each, ± SD). **E.** The luciferase reporter line *Promoter_CYP79B2_:LUC* shows an extensive activation of *CYP79B2* promoter by *Vl*43 (3 dpi) compared to mock. Upper panel: black/white pictures of the system. Lower panel: intensities of luminescence are given in false colours from low (blue) to high (red). **F.** Induction of marker genes in WT leaves at the indicated time points. Values of *Vl*43 infected samples are given relative to mock samples (set to 1) (*n* = 3 each, ± SD). **G.** Amount of *Vl*43 fungal DNA in WT, *cyp79b2/b3* and *ERF105 OE* as quantified in leaves, 2 dpi. (*n* = 3 each, ± SD). Fig.1 statistics: student’s *t*-test relative to mock (D, F) or WT (G), *p ≤* 0.05, ** *p ≤* 0.01, *** *p ≤* 0.001.

### The soil-based system is well suited for susceptibility assays

Since Koike *et al*. (1994) published a method for successfully initiating *Verticillium* disease in the greenhouse by dipping roots into spore suspensions, this protocol has been modified frequently, including the optimizations in this work. Here, approximately one week after root inoculation, fungal DNA was detectable in *Arabidopsis* leaves and its amount increased over time (Fig. 2A). From 21 dpi on, leaves turned yellow and other symptoms such as stunting (reduced rosette size) and wilting developed (Fig. 2B, Appendix S2B, C). Due to the development of disease symptoms, green leave area and biomass (fresh weight) of plant rosettes were significantly lower in infected WT than in mock treated WT at 21 and 28 dpi (Fig. 2C). Interestingly, when analysing gene expression in leaves, IG markers were not as much up-regulated as in the other infection systems and they were just slightly induced at the beginning and the end of the time course (Fig. 2D). This might be due to IG being more important in roots than in leaves during defence of *V. longisporum*. However, relative fresh weight from infected *cyp79b2/b3* plants was significantly less than from infected WT (Appendix S2D), although the difference was not dramatic. Stronger effects were obtained through quantification of fungal DNA. In line with the results above, the *cyp79b2/b3* knockout mutant had significantly more and the *ERF105 OE* significantly less fungal DNA in rosette leaves compared to WT (Fig. 2E). An alternative procedure to categorize susceptibility is quantifying fungal outgrowth from stem segments cut from infected plants. In agreement with the other results, there was more fungal outgrowth from *cyp79b2/b3* stems than from WT stems (Fig. 2F). A control experiment using *Vl*-sGFP verified that it is indeed *V. longisporum* growing out of the stems (Fig. 2G). Taken together, this system works well for susceptibility testing using both symptom development and fungal DNA amount as parameters.

**Fig. 2:**
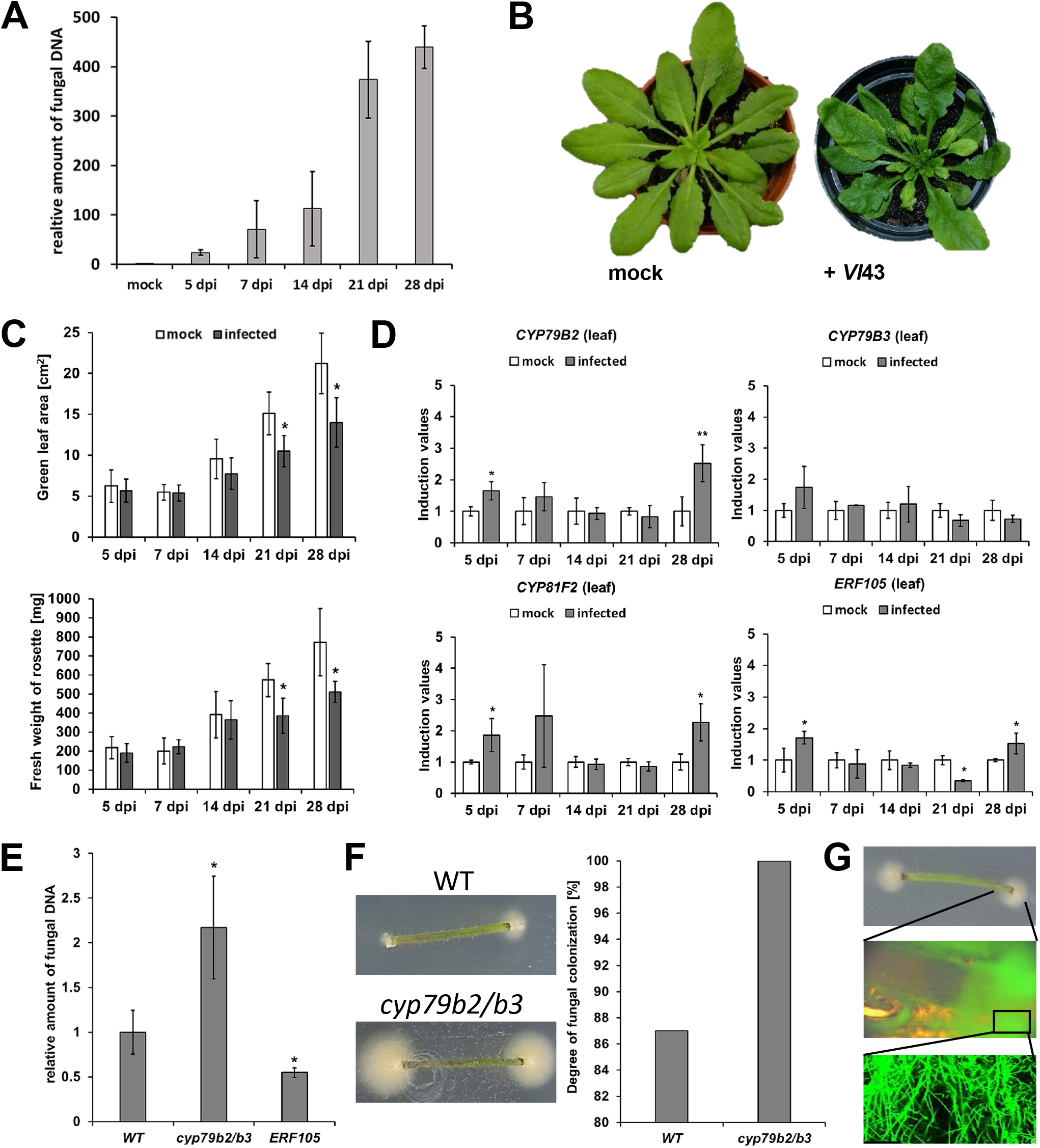
The soil-based infection system to study fungal propagation and leave responses. **A.** Relative amount of *Vl*43 fungal DNA as quantified in WT leaves at the time points indicated. Values of infected samples are given relative to background noise in mock sample (set to 1) (*n* = 3 each, ± SD). **B.** Comparison of *Vl*43 infected and mock treated plants, 21 dpi. **C.** Development of disease symptoms as quantified by green leaf area (upper part) and biomass (fresh weight, lower part) for whole rosettes of mock treated and *Vl*43 infected WT plants (*n* = 8 each, ± SD). **D.** Induction of marker genes in *Arabidopsis* WT leaves at the indicated time points. Values of *Vl*43 infected samples are given relative to mock samples (set to 1) (*n* = 3 each, ± SD). **E.** Amount of *Vl*43 fungal DNA in WT, *cyp79b2/b3* and *ERF105 OE* as quantified in leaves, 21 dpi. Values are given relative to WT (set to 1) (*n* = 3 each, ± SD). **F.** Analysis of fungal colonization in stems of *Arabidopsis*, 21 dpi: While 100 % of the stem segments from *cyp79b2/b3* were heavily colonized with *V. longisporum*, just 87 % of the WT stems were (100 stem segments tested in each case). Stem segments of *ERF105 OE* are less colonized than WT as recently published (Fröschel *et al*., 2019). **G.** Confirmation of *V. longisporum* outgrowth by using *Vl*-sGFP: stem segment with outgrowth (top), enlargement of stem end and outgrowth under fluorescence excitation (middle) and enlargement of the fluorescent hyphae (below)(adapted from Heeg, 2017). Fig. 2 statistics: student’s *t*-test relative to mock (C, D) or WT (E), *p ≤* 0.05, ** *p ≤* 0.01, *** *p ≤* 0.001.

### A novel set-up with plastic cups closes experimental gaps of other systems

Although the petri dish system is well applicable for assessing differentially expressed genes, it has weaknesses with pathogenicity assays and vice versa with the soil-based system. Therefore, we established a new system combining both susceptibility assays and expressional analyses. Reliability was assessed by comparing its results with the two existing systems. Fig. 3A briefly summarizes the main features with a lower plastic cup containing the plants in agar medium and an upper inverted cup as lid. Four plants were cultivated per cup and inoculated with *V. longisporum* in an infection channel located in the middle of the agar medium. Nicely, the fungus grew towards the roots so that a uniform infection of all plants was achieved. A time course corroborated that the fungus reached the foliage of *Arabidopsis* after inoculating at the roots (Fig. 3B). Fungal DNA was first detectable 4 dpi (not shown) and extremely increased until 16 dpi. At later time points, the curve flattened (not shown). Subtle disease symptoms occurred on leaves, such as turning yellow and reduction in rosette size (Fig. 3C). Markers for IG synthesis were transcriptionally induced in roots, as demonstrated in a time-course experiment: after reaching a maximum at 10-12 dpi, expression decreased again (Fig. 3D). This observation was similar to the petri dish system but time delayed. By comparing WT with the highly susceptible *cyp79b2/b3* mutant, it turned out that 10 dpi was too early to find significant differences. This changed at 12 dpi and 14 dpi where significantly more (4 to 6-fold) fungal DNA was observed in the mutant (Fig. 3E). In contrast, *ERF105 OE* possesses increased IG marker gene expression (Fröschel *et al*., 2019) and was more tolerant than WT, since less fungal DNA and more relative fresh weight were found in the overexpressor upon infection (Fig. 3E, Appendix S3I). Transcriptional induction of IG markers was observed in leaves, too. Maxima occurred at the beginning and at the end of the time-course (Fig. 3F), a pattern similar to that in the soil-based system.

**Fig. 3:**
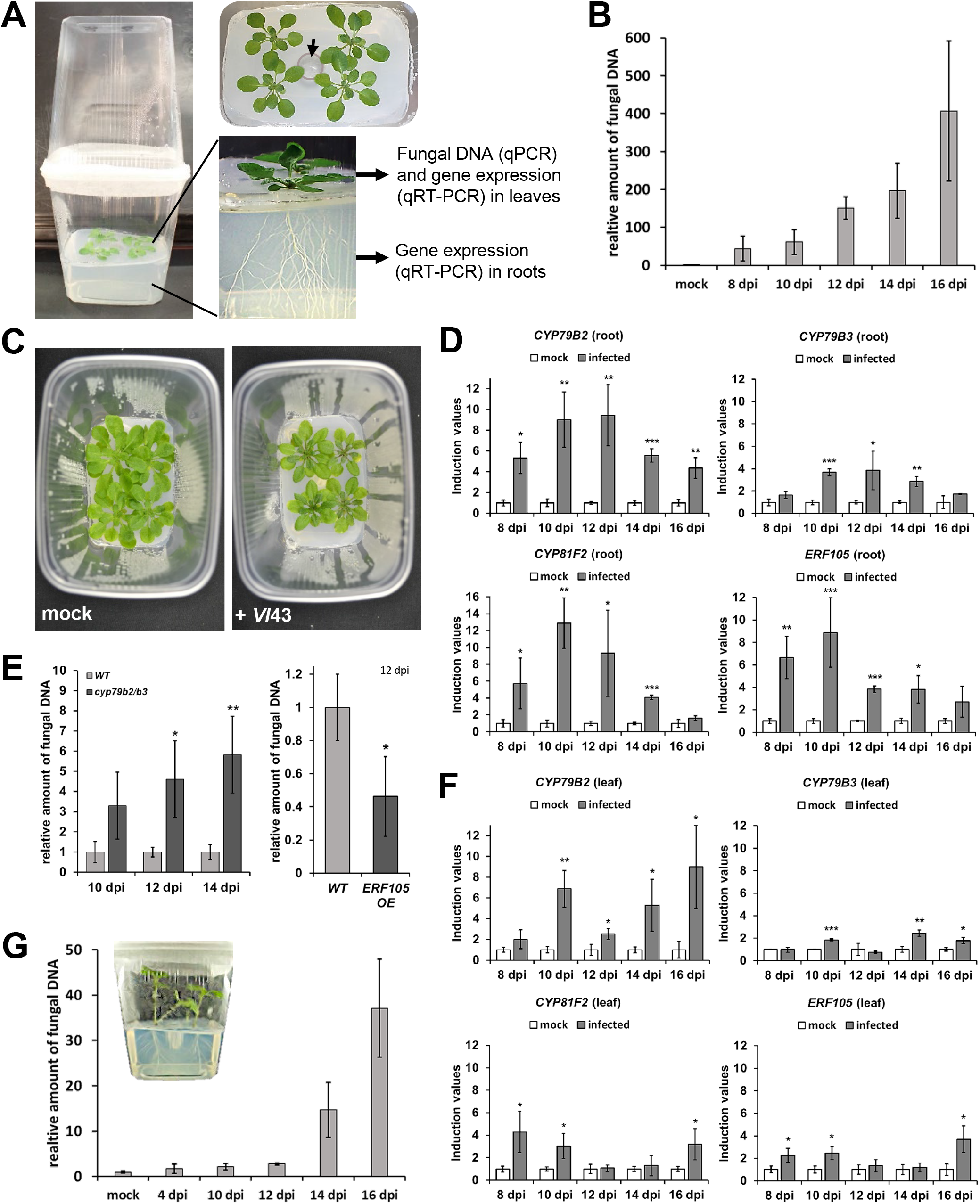
The *in vitro* infection system in plastic cups discloses root and leave responses as well as fungal propagation. **A.** Basic set-up of the system: four plants grow on agar medium in the lower plastic cup, while an inverted cup serves as lid. Conidiospores are added in a channel in the middle of the plants (black arrow). A separating plastic layer on the agar medium prevents a direct contact of the leaves with spreading *V. longisporum*. Therefore, all fungus detectable in leaves results from *in planta* spread. Possibilities for downstream analyses are given. **B.** Relative amount of *Vl*43 fungal DNA in *Arabidopsis* WT leaves at the time points indicated. Values of infected samples are given relative to background noise in mock sample (set to 1) (*n* = 3 each, ± SD). **C.** Representative photos from mock treated and *Vl*43 infected *Arabidopsis* WT plants, 12 dpi. **D.** Induction of marker genes in *Arabidopsis* WT roots at the indicated time points. Values of *Vl*43 infected samples are given relative to mock samples (set to 1) (*n* = 3 each, ± SD). **E.** Amount of *Vl*43 fungal DNA as quantified in *Arabidopsis* leaves: in WT and *cyp79b2/b3* at the time points indicated (left) and in WT and *ERF105 OE*, 12 dpi (right). All values are given relative to WT (set to 1) (*n* = 3 each, ± SD). **F.** Induction of marker genes in *Arabidopsis* WT leaves at the indicated time points. Values of *Vl*43 infected samples are given relative to mock samples (set to 1) (*n* = 3 each, ± SD). **G.** Relative amount of *Vl*43 fungal DNA as quantified in *Brassica napus* WT leaves at the time points indicated. Values of infected samples are given relative to background noise in mock sample (set to 1) (*n* = 3 each, ± SD). Inlay: cup with *B. napus*. Fig.3 statistics: student’s *t*-test relative to mock (D, F) or WT (E), *p* ≤ 0.05, ** *p* ≤ 0.01, *** *p* ≤ 0.001.

When *Brassica napus* plantlets were root inoculated, the amount of fungal DNA in leaves also increased with time (Fig. 3G). While fungal propagation was slow in the beginning, it rapidly increased from 12 dpi on. In a nutshell, the novel infection system in plastic cups is well applicable and reliable for studying *V. longisporum* disease in *Arabidopsis* and also in the agronomically interesting model plant *Brassica napus*.

## DISCUSSION

This study revealed advantages and disadvantages of three versatile infection systems by using selected examples with the model root pathogen *Verticillium longisporum*. This aimed at assisting researchers to find an appropriate infection system for their specific research question.

### Advantages and disadvantages of the infection system in petri dishes

It is a quick set-up leading to results within 2 weeks. During the whole procedure, the system can be kept sterile, which prevents disturbing contaminations.

Immediately after adding the spores, they are in close contact with the roots allowing investigations of early responses. Compared with the other two infection systems, transcriptional responses are very vigorous in leaves and roots, which might be considered artificial in terms of strength as the plantlets are relatively small and quickly overcome by pathogens. Nevertheless, as shown with the transcriptional induction of IG markers, expressional changes can be studied in both roots and leaves. A great advantage here is that many plants are used in pools (approx. 100 per plate), which reduces fluctuations due to variations in infection usually observed in plant pathogen interactions (Kosman *et al*., 2019). Genome-wide −omics approaches are feasible such as Microarray, RNA-sequencing or even cell-layer specific assays (Iven *et al*., 2012; Fröschel *et al*., 2020). Expressional changes in roots can be easily corroborated with adequate reporter-lines (e.g., GFP, GUS or luciferase) and dynamics of fungal spread can be visualized with the GFP-tagged *V. longisporum* strain. Root phenotypes are well observable and quantifiable through measuring primary root growth or lateral root development upon microbial challenge. Using the model organisms *V. longisporum* and *A. thaliana*, the petri dish system is not suitable to investigate time-points upon 6 dpi as plants are progressively harmed and dying. Additionally, the medium dries out over time. Susceptibility tests at relatively early time points (1 to 2 dpi) are practicable as long as differences regarding WT are substantial as demonstrated for the *cyb79b2/b3* double mutant. Subtle differences in susceptibility are difficult to detect with this system. In this case, it is recommended to choose one of the other two infection systems as they are more sensitive in analysing fungal spread, like demonstrated with the tolerant *ERF105 OE*. Quantification of general disease symptoms, like reduction in biomass, is hardly practicable in petri dishes and you rely on the measurement of fungal DNA as the best parameter. Since the system is small, it is hardly usable for larger *B. napus* seedlings. Recently, Behrens *et al*. (2019) established another infection system on plates, where a brush was dipped in *V. longisporum* conidia suspension and used to distribute spores along roots growing onto an agar medium. This method might be considered as alternative.

### Advantages and disadvantages of the infection system in plastic cups

A yet unpublished *in vitro* infection system in plastic cups was established to investigate infection time points located between the petri dish system (hours post inoculation to 6 dpi) and the soil-based system (> 21 dpi). This novel system is easy to handle, not very space consuming and it can be kept germ-free from external contaminations, so that bilateral interactions remain undisturbed. In total, the experimental setup needs 4-5 weeks to complete with results.

Expressional changes can be analysed in roots and shoots leading to results comparable to the petri dish system, although with lower induction values for some markers. Nevertheless, since roots are embedded in agar medium and not as easy to access as in petri dishes, confirmation of expressional changes in roots with reporter lines are more difficult here (pulling out the roots leads to injury, which might interfere with reporter gene expression). Progression of infection is slower compared to the petri dish system reflecting more natural conditions. Most suitable time points to perform pathogenicity tests with different *Arabidopsis* lines range between 10 and 16 dpi. Much later time points with *Arabidopsis* are not recommendable, as the curve of fungal growth reaches a plateau. From then on, the plants might be extensively colonized and overcome, which leads to difficulties in recognizing differences between lines. However, the best time-point should be defined through preliminary experiments, as it was exemplified with *cyp79b2/b3* showing significant differences compared to WT from 12 dpi on. A loss-of-function in the IG pathway increases susceptibility, while a gain-of-function enhances tolerance, which is in line with the other experiments in this study and previous publications (Iven *et al*., 2012; Fröschel *et al*., 2019). This confirms the reliability of the new system in plastic cups and its applicability to analyse fungal propagation in different transgenic lines. Therefore, identification of differentially expressed genes in roots and shoots as well as mutant analysis are doable in parallel underlining the substantial advantage of this system. Regarding *B. napus*, this infection system is also suited for the larger seedlings, but later time points (14 to 20 dpi) are advisable for susceptibility tests, as fungal spread is delayed and weaker in comparison to *Arabidopsis*. Other brassicaceous model plants could also be applied there (e.g., broccoli).

Interestingly, *V. longisporum* reaches the foliage after inoculating at the roots without damaging root tissues. qPCR analysis demonstrated a rapid colonization of the foliage in the agar-based *in vitro* systems (petri dishes, plastic cups), which reflects propagation via the xylem. In petri dishes, this is a very rapid process within 24 hpi. As the endodermis provides a barrier to restrict *V. longisporum* propagation to the vasculature (Fröschel *et al*., 2020), the fungus might penetrate either through not fully developed tissues (e.g., meristematic zones) or at sites, where lateral roots perforate outer cell-layers (Yadeta and Thomma, 2013) or it weakens the barrier actively. This observation in the agar-based infection systems is new, since it was anticipated that *Arabidopsis* roots need to be wounded to facilitate *V. longisporum* to enter, like it is the case in the soil-based infection system.

### Advantages and disadvantages of the soil-based infection system

Since roots are damaged due to uprooting and washing prior dipping, this opens a gate for *V. longisporum* to enter the vasculature directly, which makes this system a bit artificial. On the other hand, it could mimic natural conditions that injure roots, such as nematode feed (e.g., Back *et al*., 2002). However, of the three systems discussed, it is the one closest to natural conditions as plants grow in soil. The infection progression is slower compared to the other two systems and symptom development can be tracked (reduction in biomass, green leaf area). Analysing colonization in stems is well suited to specify increased tolerance or susceptibility in extensive screenings of ecotypes in genome-wide association studies (GWAS) or of transgenic plant collections (e.g., *At*TORF-Ex, Fröschel *et al*., 2019). Since infected plants are smaller than mock treated ones, the measurement of height might be an additional parameter to quantify symptoms (Fröschel *et al*., 2019). Nevertheless, symptom quantification can lead to huge error bars due to individual variations. Symptom-derived results should always be validated by quantifying fungal DNA in leaves, which is by far the most reliable method. With *Arabidopsis*, best results are achieved here at 21 dpi.

Applying this system, it takes 6-7 weeks to get results, it needs more space than the others and the conditions are considered to be semi-sterile only so that the bilateral interaction is not completely undisturbed. Pathogen-triggered expressional changes can be investigated in leaves, but hardly in roots. Since roots grow in soil, it is difficult to clean them from soil sufficient for qRT-PCR analysis without the risk of reprogramming gene expression due to washing. This is a limitation, at least for the markers discussed. The response to *V. longisporum* was weak in leaves and, from the experience with the other two systems, roots appear to respond stronger.

Plantlets from *B. napus* can be successfully inoculated with this method similar to *Arabidopsis* as previously described by Singh *et al*. (2012). They infected 7 d old *B. napus* seedlings by up-rooting and subsequent root dip inoculation, and found that the highest concentration of fungal DNA was present in hypocotyl stems. Thus, quantification of fungal DNA in *B. napus* seems to be more reliable with hypocotyls than with leaves (> 28 dpi).

### Conclusions

Expressional induction of the IG pathway is one important marker to evaluate *V. longisporum* infection. It is more central in roots than in shoots and more dominant in the beginning of infection. Assays with mutants and overexpressors in the IG pathway demonstrate that these are important secondary metabolites involved in the restriction of *V. longisporum* (Iven *et al*., 2012, Fröschel *et al*., 2019). The given examples highlight the ability of the three described infection systems to further decipher root-microbe interactions, each with its specific pros and cons. Although the focus of this work was set on *V. longisporum*, other soil-borne microorganisms can be used in the described infection systems, too. Hence, very recently we applied the soil-borne, beneficial fungus *Serendipita indica* successfully in the petri dish system to examine differentially expressed genes in roots (Fröschel *et al*., 2020). Additionally, the novel *in vitro* infection system in plastic cups might be feasible for studies with *S. indica* or even with non-fungal microbes such as soil-borne Ooymcetes (e.g., *Phytophthora parasitica*), at least some preliminary tests speak for it.

## ACKNOWLEDGEMENTS

I would like to thank Alexander Marsell (University of Würzburg), who contributed to the study as a highly motivated student, and Jaqueline Komorek for their assistance. Furthermore, I acknowledge Wolfgang Dröge-Laser, Frank Waller and Christoph Weiste (all at University of Würzburg) for supporting this study and for rewarding proof-reading.

## Author contributions

C.F. established the novel infection system in plastic cups, performed all the experiments in this study and wrote the manuscript.

