## Appendix S1 for "In-depth evaluation of root infection systems with the vascular fungus *Verticillium longisporum* as soil-borne model pathogen"

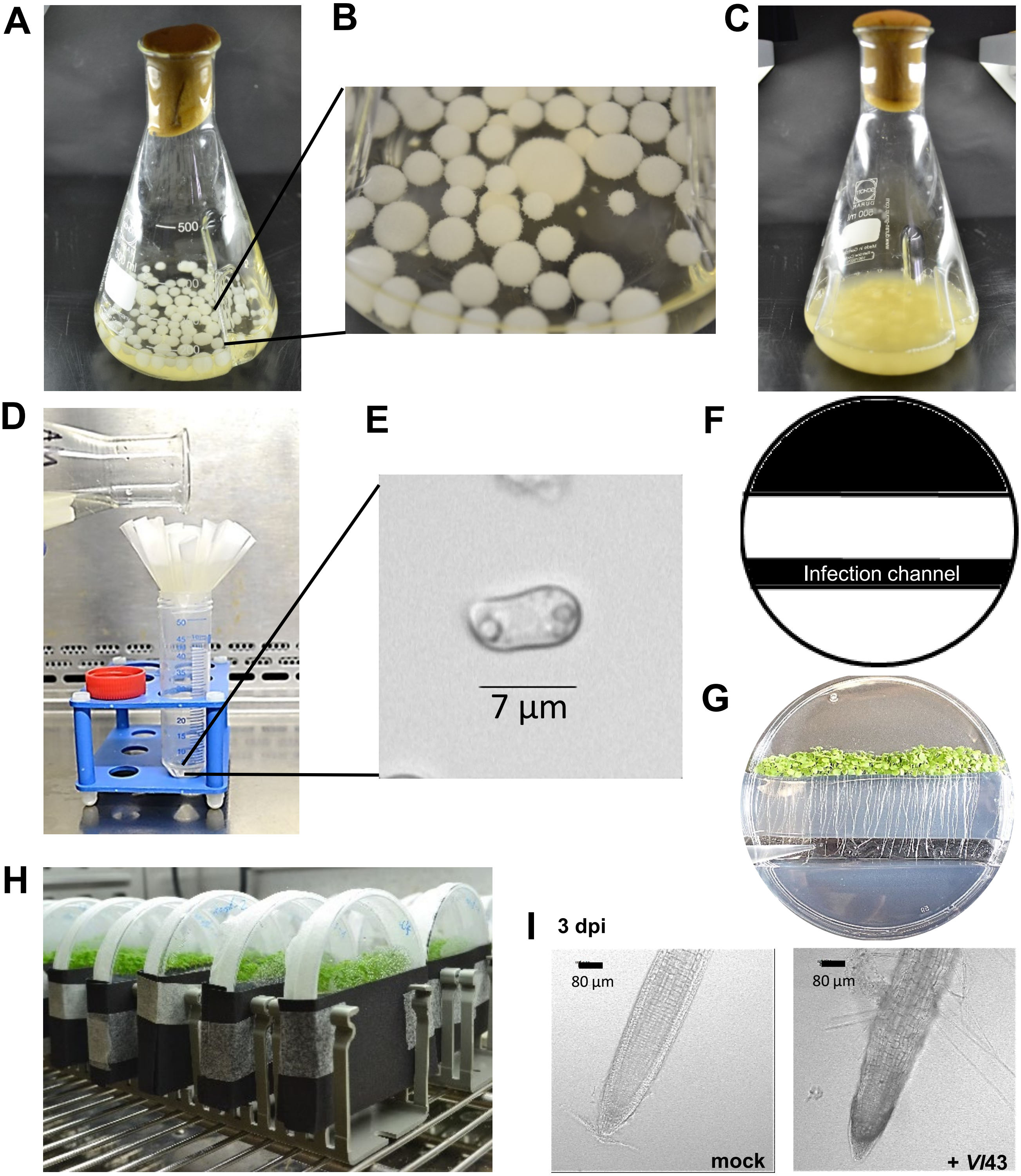


**Appendix S1: Preperation of *Verticillium longisporum* inoculum and infection system in petri dishes.** All steps should be carried out in a laminar flow hood with sterile equipment. **A.** and **B.** Spores from a long time storage were transferred to PDB liquid medium and incubated under continuous shaking. 7-10 days later, white mycelium balls had emerged. **C.** 4-5 days after the PDB medium had been replaced by liquid CDB medium, the supernatant became yellow-greyish due to the formation of conidiospores. **D.** Spores were purified through folded filter paper into a 50 ml tube. **E.** Characteristic long-drawn conidiospore under the microscope. **F.** Black areas in the scheme were removed from the solidified agar-medium with a scalpel. **G.** 50-100 *Arabidopsis* plantlets were cultivated per plate and roots were inoculated via adding the spore solution into the infection channel. **H.** The Leukopor® sealed plates were set up vertically and covered with black papers at the roots. This allowed darkened conditions to roots and soil-borne fungus. **I.** Infected roots (+ *Vl*43) were different from mock treated ones as it was observable under the microscope (3 dpi).
