## Appendix S2 for "In-depth evaluation of root infection systems with the vascular fungus *Verticillium longisporum* as soil-borne model pathogen"

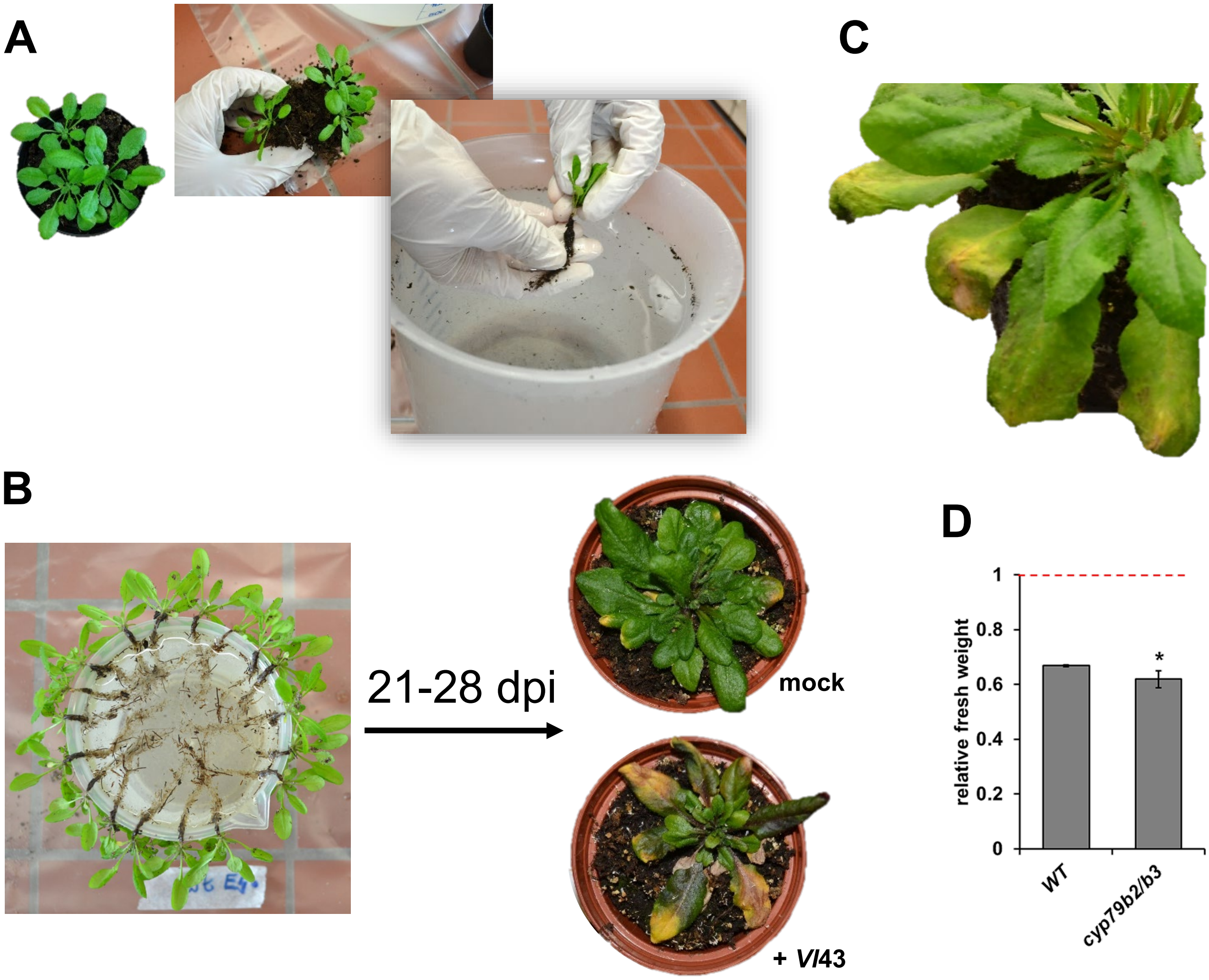


**Appendix S2: Overview of the soil-based infection system in pots with *Arabidopsis*. A.** 3-4 plants were cultivated per pot in a soil-sand mixture. To moderate phenotypical and individual variations that might influence experimental outcome, plenty of plants were cultivated and just plants of similar size were carefully selected for the experiments. 21 d old plants were prepared for “root dip inoculation”: Carefully turning the pots without destroying the rosettes, excavating of single plants and washing of roots in a 5-liter jar. Only roots were gently washed with one hand, while the rosettes were kept out of the water with the other hand. In general, this procedure injures roots, making it easier for *V. longisporum* to penetrate them. **B.** All plantlets were prepared and temporarily stored hanging on a small cup with the roots in water and the rosettes outside. After all plants were prepared like this, they were rapidly transferred into petri dishes containing the spore solution or mock. This ensured the same starting point of inoculation for all plants. Like in the preparation shown in the photo, just the roots were hanging in the spore solution, while the rosettes did not had contact to the solution. After 60 min incubation, plants were repotted into single pots containing soil. 21-28 dpi, observable symptoms developed in infected plants compared to mock treated ones (demonstrated in the photos with *Arabidopsis* WT plants, 28 dpi). **C.** Enlarged section of wilted and yellowish leaves from infected *Arabidopsis* WT, which can be described as typical disease symptoms. **D.** Differences in infection progression between plant lines can be quantified either by fungal DNA measurement like it is exemplified in the main figures or via development of symptoms. Here, relative fresh weight of WT and *cyp79b2/b3* was determined, 21 dpi. Fresh weight of mock controls from both lines was set to 1 (red dashed line) and infected samples are given relative to mock. Both, infected WT and *cyp79b2/b3* plants showed reduced fresh weight compared to their mock. Relative fresh weight of *cyp79b2/b3* was even more reduced than the one of WT ($n$ = 15 each, ± SEM, student‘s $t$-test relative to WT, * $p \leq 0.05$).
