## Appendix S3 for "In-depth evaluation of root infection systems with the vascular fungus *Verticillium longisporum* as soil-borne model pathogen"

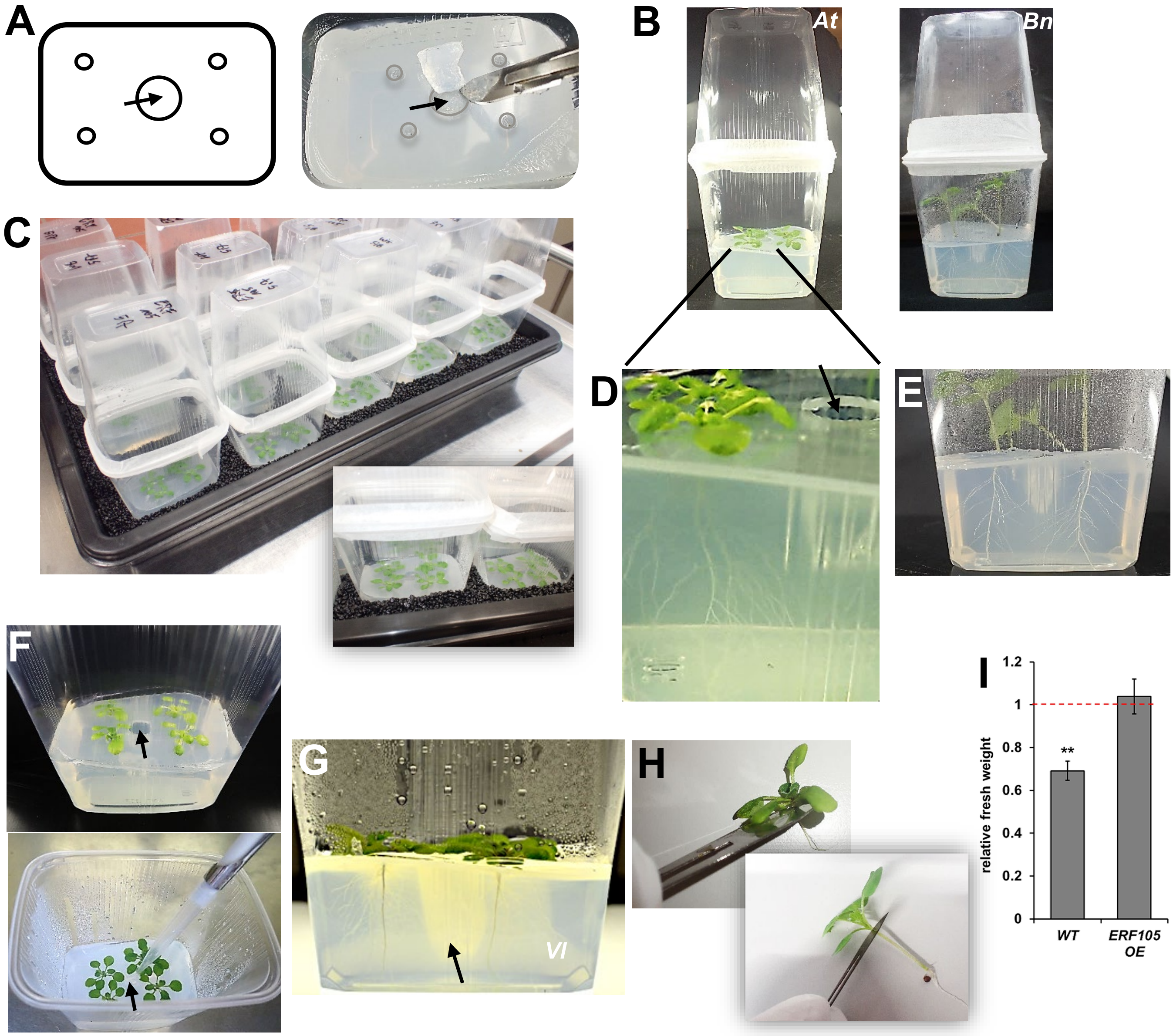


**Appendix S3: Overview of the sterile *in vitro* infection system in plastic cups.** All steps should be carried out in a laminar flow hood with sterile equipment. In all pictures, the arrows indicate the infection channel. **A.** Separating plastic layer as scheme (left) to illustrate the prefabricated holes and as photo (right) with the cut of the infection channel in the centre and the four smaller holes for the seeds. **B.** The lower plastic cup with the plants growing on agar medium was covered with a second and inverted cup as lid. Closure with Leukopor® allowed gas exchange. Medium for *Arabidopsis* (*At*, left) filled approx. ⅓ of the lower cup, while for *Brassica napus* (*Bn*, right) more medium was needed (approx. ½ - ⅔ of the lower cup). **C.** To darken the roots, the lower parts of the cups were embedded in granulate (black split, 2-3 mm). **D.** Enlarged section with *Arabidopsis*. The separating layer can be seen that prevents a direct contact of the leaves with the fungus after root inoculation. Therefore, all fungal DNA detectable in leaves derived solely from fungal spread from root to shoot within the plant. **E.** Enlarged section with *Brassica napus*. **F.** Inoculation of 21 d old *Arabidopsis* seedlings: Adding a solution containing *V. longisporum* conidiospores into the infection channel. **G.** Several days after inoculation, mycelium (*Vl*) became visible in the medium. **H.** From *Arabidopsis*, the whole rosettes were harvested with a scalpel. Roots were carefully pulled out of the medium and blotted with paper towel to remove agar remains. From *Brassica napus*, all leaves were harvested but not the stems. **I.** Differences in infection progression between plant lines can be quantified either by fungal DNA measurement like it is exemplified in the main figures or via development of symptoms. Here, relative fresh weight of WT and *ERF105 OE* was determined, 12 dpi. Fresh weight of mock controls from both lines was set to 1 (red dashed line) and infected samples are given relative to their mock. While infected *ERF105 OE* plants showed almost mock-like fresh weight, fresh weight of infected WT significantly decreased ($n$ = 15 each, ± SEM, student's $t$-test relative to mock, ** $p \leq 0.01$).
