## Appendix S4 for "In-depth evaluation of root infection systems with the vascular fungus *Verticillium longisporum* as soil-borne model pathogen"

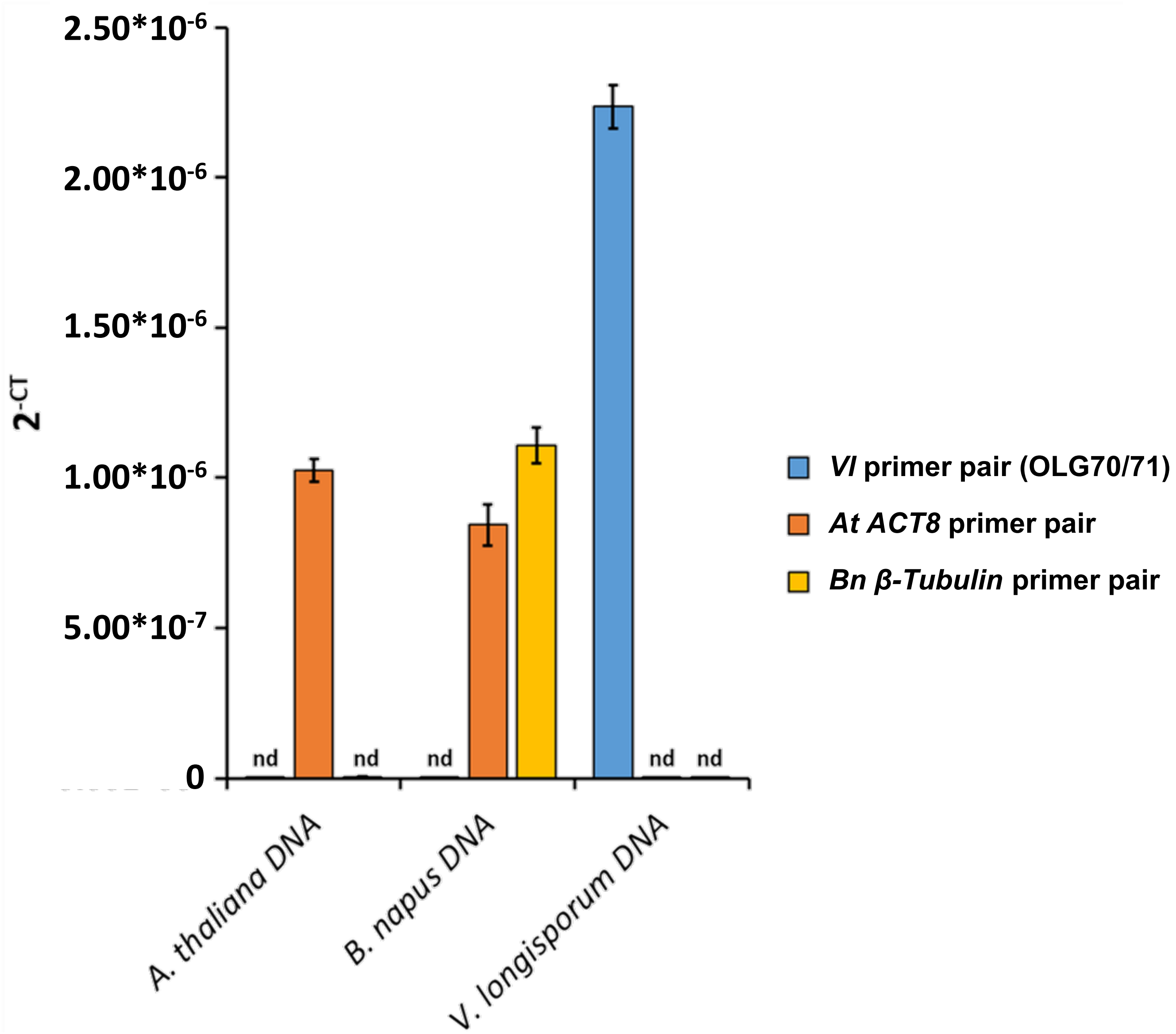


**Appendix S4: Analysis of primer specificity for the organisms under investigation.** Genomic DNA was isolated from either uninfected *A. thaliana* (*At*) or *B. napus* (*Bn*) leaves. *V. longisporum* genomic DNA was isolated from freshly harvested conidiospores. 100 ng of each genomic DNA extract served as template in qPCR with either *At ACT8*, *Bn ß-Tubulin* or *Vl* specific primers. $2^{-CT}$ values are given on y-axis ($n$ = 3 each, ± SD). *At ACT8* primers led to amplification in DNA extracts of the two related species *A. thaliana* or *B. napus*, but not with *V. longisporum* DNA. Therefore, this primer pair can be used for normalization in infection experiments with *A. thaliana* and *B. napus*, respectively. *Bn ß-Tubulin* primers led specifically to an amplicon in *B. napus* DNA extract. The *Vl* primer pair OLG70 / OLG71 amplifies a specific *Verticillium* fragment encoding parts of the 5S rRNA (Eynck *et al*., 2007; Singh *et al*., 2012). The signal with the *Vl* primer pair was high with *V. longisporum* DNA extract as template, but almost not detectable with *A. thaliana* or *B. napus* DNA, although there was a bit of background noise interpretable as unspecific binding of the primers. This background noise with *Vl* primers in uninfected plant samples (mock) was sufficient enough to use it in the time course experiments to calculate the mock value and to normalize the infected samples to mock; nd = not detectable (defined as values $< 3*10^{-10}$).
