## Appendix S5 for "In-depth evaluation of root infection systems with the vascular fungus *Verticillium longisporum* as soil-borne model pathogen"

**Appendix S5: Primer-Oligomers used in this study**

Primer sequences for qRT-PCR and qPCR:

| Gene ID | Primer Name | Sequence (5'->3') |
|---|---|---|
| AT4G39950 | *CYP79B2* for | GAGACGACGCCGTATATATTTTATG |
| AT4G39950 | *CYP79B2* rev | GGAGACACAATTCAAGTGTCCCAA |
| AT2G22330 | *CYP79B3* for | ACCGGAAAGAGAGGATGTGCTG |
| AT2G22330 | *CYP79B3* rev | CGCTAGCATCATGGTCGTTATCGC |
| AT5G57220 | *CYP81F2* for | GAAGATGTTGACATGACAGAG |
| AT5G57220 | *CYP81F2* rev | TGCTTAAACCGGTAAACTTC |
| AT5G51190 | *ERF105* for | AGCTGCAAGAGGTTATGAC |
| AT5G51190 | *ERF105* rev | TTCATCCTCCGTCACCTGTC |
| AT3G62250 | *UBQ5* for | GACGCTTCATCTCGTCC |
| AT3G62250 | *UBQ5* rev | GTAAACGTAGGTGAGTCCA |
| AT1G49240 | *At ACT8* for | GGTTTTCCCCAGTGTTGTTG |
| AT1G49240 | *At ACT8* rev | CTCCATGTCATCCCAGTTGC |
| *B. napus* | *Bn* β-Tubulin for | AGGTCTCCGACACTGTTGTTG |
| *B. napus* | *Bn* β-Tubulin rev | GGAGTTGAGTTGACCAGGGA |
| *Verticillium 5S rRNA* | *OLG70* | CAGCGAAACGCGATATGTAG |
| *Verticillium 5S rRNA* | *OLG71* | GGCTTGTAGGGGGTTTAGA |
