## Appendix S6 for "In-depth evaluation of root infection systems with the vascular fungus *Verticillium longisporum* as soil-borne model pathogen"

A

B

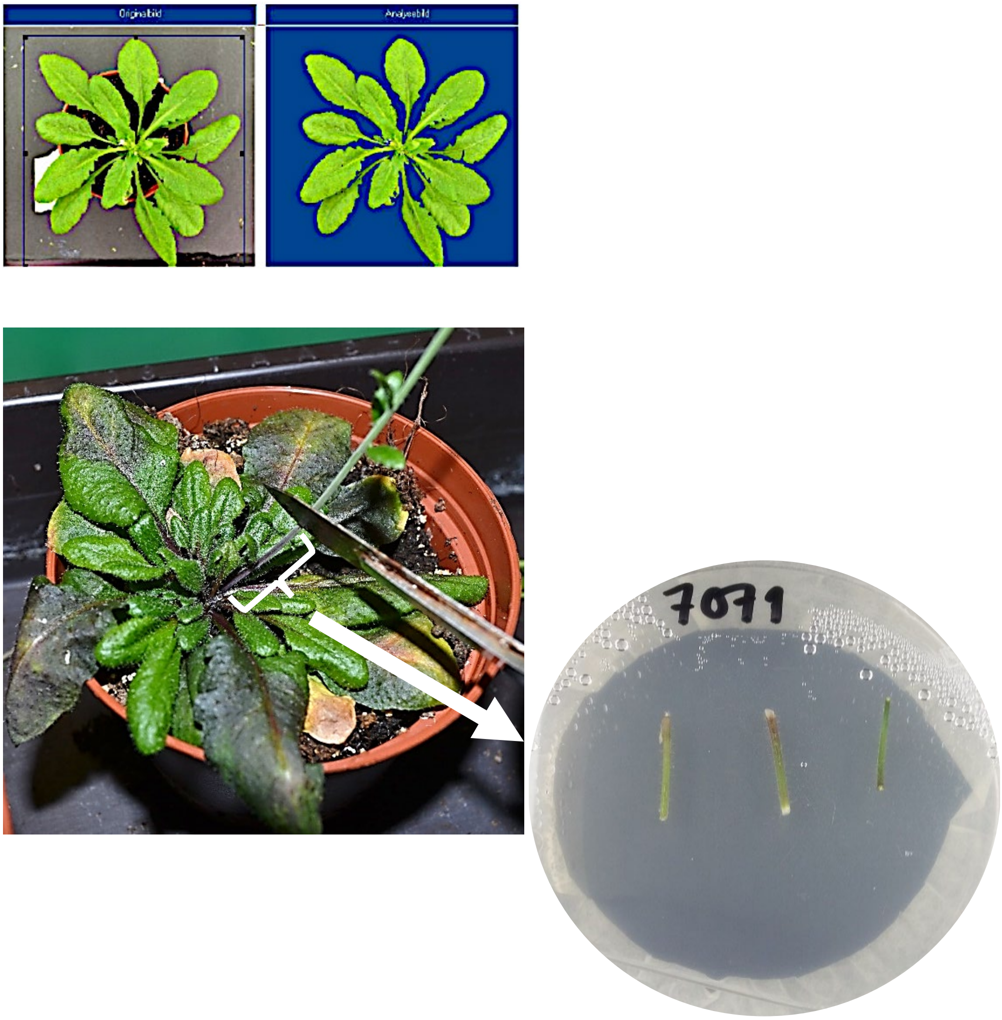


**Appendix S6: Methodological details presented for better understanding. A.** Infection symptoms were monitored by measuring the green leaf area on digital photos. The projected leaf area of the complete rosette was determined with the image analysis program BlattFlaeche® (Datinf GmbH, Tübingen, Germany). The settings were chosen so that only the green area was measured: the original picture on the left and the projected green leaf area on the right. If yellow leaf areas occurred, they were not projected. **B.** 1–1.5 cm of inflorescence stem segments were cut from the bottom, surface sterilized and placed on solidified Potato Dextrose Broth Agar. After 3-5 d, fungal outgrowth occurred, which could be categorized.
